# Sesamol and its derivative investigated as antiandrogen - A potential prevention to prostate cancer in rats

**DOI:** 10.1101/2021.04.22.440973

**Authors:** Abhishek Shah, Aarti Abhishek Shah, Deelip S. Rekunge, Aravinda Pai, Ganesh U. Chaturbhuj, K Nandakumar, Richard Lobo

## Abstract

Androgen signaling is essential for the development of prostate cancer (PCa) initiated from prostatic basal cells with collocation of androgen receptor gene mutations. Phytoestrogens, the naturally occurring compounds are AR antagonist. These compounds downregulate prostate-specific antigen (PSA) expression and cell proliferation. Thus, this gives a track to research these compounds as a possible treatment for PCa. In this work, STITCH and molecular docking predict the conformation of ligands inside the suitable target binding site. Therefore, a study was planned to know the interactions among SM and its derivatives with AR. It was further, evaluated for *in vitro* evaluation on LNCaP, PC-3, and DU-145 using MTT studies. The two lead compounds shortlisted from MTT studies were further analyzed for androgen-regulated genes by using RT-PCR, western blot studies and an animal model of prostate cancer. We found that SM and its derivative (3’-MA) may prevent the development of PCa by androgen pathway.

## 1 Introduction

Prostate cancer (PCa) is the second leading malignancy amongst men worldwide. Androgen signaling is the driving force for the progression of PCa. [1][2][3] AR regulates multiple cellular events, proliferation, apoptosis, migration, invasion, and differentiation.[4][5] According to the latest finding by Yongfeng he (2018), androgen signaling is essential for the development of PCa initiated from prostatic basal cells[6] Collocation of androgen receptor gene mutations in PCa.[7] Thus, AR is the primary drug target for PCa treatment. The androgens are key players for the growth of PCa in the initial stages, which leads to a treatment choice as androgen deprivation therapy (ADT). The AR is essential in all stages of PCa progression, it may be a relapse of the tumor, which can be even androgen-independent in nature.[8] The oral dosage form of antiandrogens are freely available and based on chemical structure, it can be classed as steroidal and non-steroidal. Cyproterone acetate being steroidal and bicalutamide, flutamide, and nilutamide are non-steroidal.[9] Although ADT is a highly effective treatment option, it is associated with considerable side effects. It includes hot flashes, insulin resistance, cardiovascular (CV) diseases, abridged body mass and muscle strength, others relating to bones osteoporosis and major concerns maybe depression and sexual dysfunction. [10]

In this context, there is a need for investigating natural antiandrogens to alleviate adverse effects arising from ADT and subsequently enhance the patient’s quality of life. The examples of natural antiandrogens are Mahanine, 2′-hydroxy genistein-7-O-glucoside, 3, 3′-Diindolylmethane, N-butyl benzene sulfonamide etc.[11] Phytoestrogens, the naturally occurring compounds are AR antagonist. These compounds downregulate the activation of PSA and cell proliferation. Thus, this gives a track to research these compounds as a possible treatment for PCa.[12][13]

Sesamolin is also a precursor to sesamol and SM is formed in frying procedures.[14] Sesame seed is a rich source of dietary lignans.[15] Lignan (sesamol, sesamin, and sesamolin) profile was identified in sesame seeds.[16][17] SM is the main reason for the oxidative stability of sesame seed oil. [18] Pianjing P, et al. investigated the phytoestrogenic potential of Sesame Lignans and their metabolites on human breast cancer cells.[19] In the present study, we claimed and tested the SM and its derivatives to be used as a prophylactic treatment for prostate cancer.

## 2 Materials and Methods

### 2.1 STITCH

Understanding the interactions between SM and androgen-regulated genes is essential to predict the molecular mechanism for the androgen signaling pathway. The online database tool “STITCH (‘search tool for interactions of chemicals’) was used to understand the interaction of Sesamol and associated genes as androgen receptor gene (AR), FKBP5 (FKBP Prolyl Isomerase 5), Prostate-specific antigen (PSA/KLK3), Transmembrane Serine Protease (TMPRSS2) was used (http://stitch.embl.de) [17]. The input and predicted functional partners are shown in table 2 and table 3 respectively.

**Table 1:**
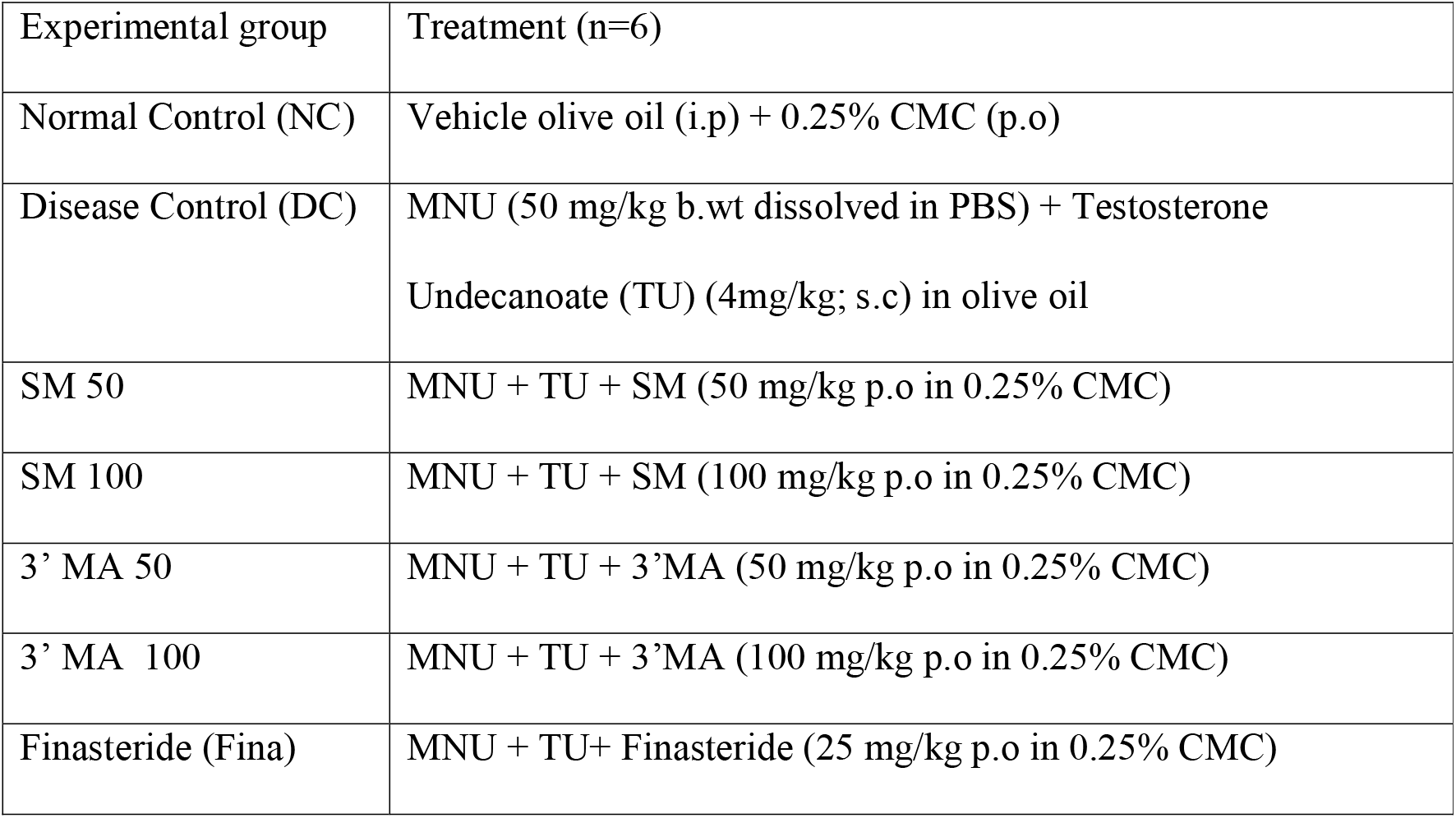
Animal grouping and treatment

| Experimental group | Treatment (n=6) |
| --- | --- |
| Normal Control (NC) | Vehicle olive oil (i.p) + 0.25% CMC (p.o) |
| Disease Control (DC) | MNU (50 mg/kg b.wt dissolved in PBS) + Testosterone<br>Undecanoate (TU) (4mg/kg; s.c) in olive oil |
| SM 50 | MNU + TU + SM (50 mg/kg p.o in 0.25% CMC) |
| SM 100 | MNU + TU + SM (100 mg/kg p.o in 0.25% CMC) |
| 3' MA 50 | MNU + TU + 3'MA (50 mg/kg p.o in 0.25% CMC) |
| 3' MA 100 | MNU + TU + 3'MA (100 mg/kg p.o in 0.25% CMC) |
| Finasteride (Fina) | MNU + TU+ Finasteride (25 mg/kg p.o in 0.25% CMC) |

**Table 2:** Input for STITCH analysis

|  |  |  |  |
| --- | --- | --- | --- |
| Sesamol | Lignans | Phytoestrogens | AR |

**Table 3:**
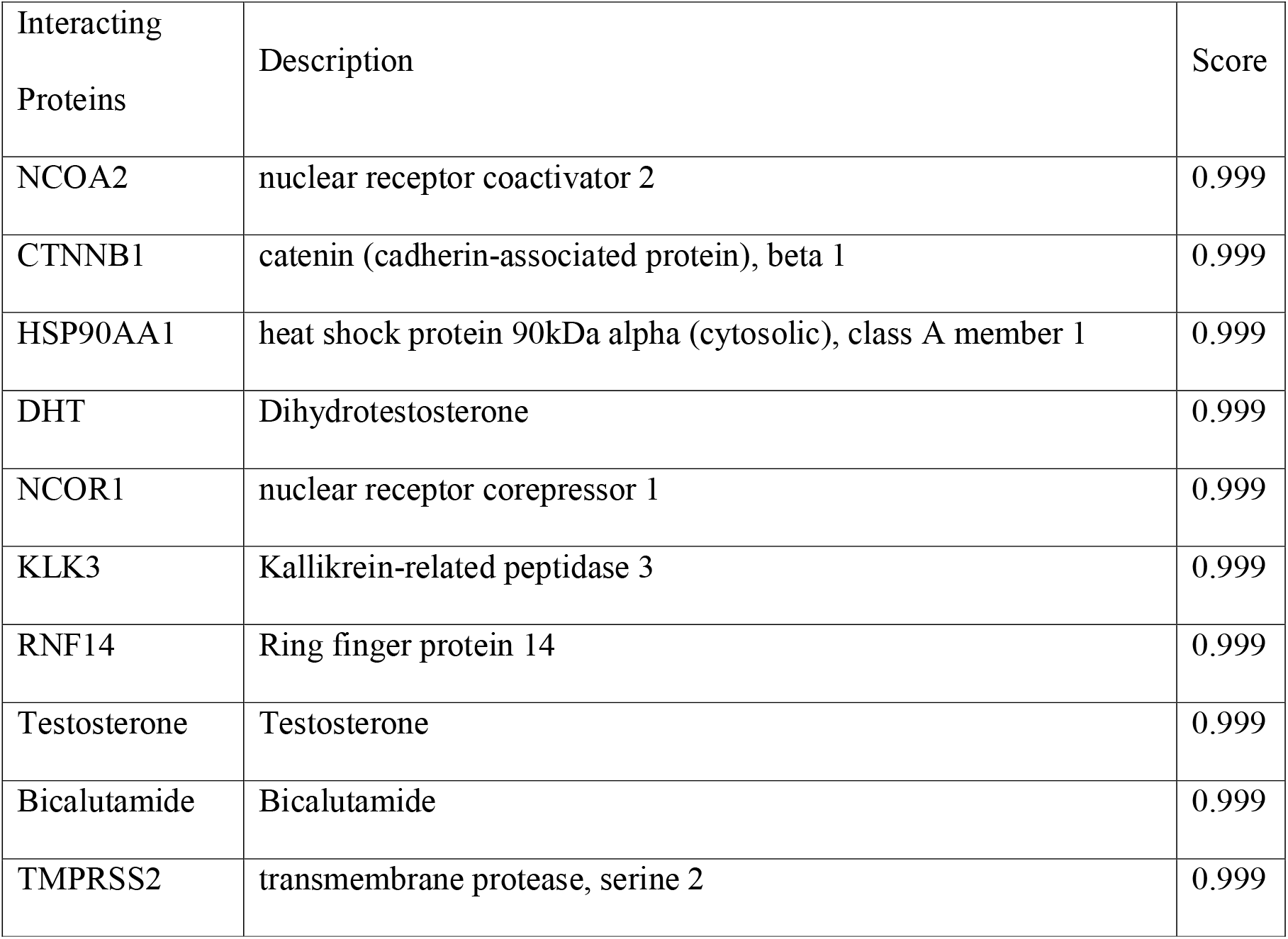
Predicted Functional Partner

| Interacting<br>Proteins | Description | Score |
| --- | --- | --- |
| NCOA2 | nuclear receptor coactivator 2 | 0.999 |
| CTNNB1 | catenin (cadherin-associated protein), beta 1 | 0.999 |
| HSP90AA1 | heat shock protein 90kDa alpha (cytosolic), class A member 1 | 0.999 |
| DHT | Dihydrotestosterone | 0.999 |
| NCOR1 | nuclear receptor corepressor 1 | 0.999 |
| KLK3 | Kallikrein-related peptidase 3 | 0.999 |
| RNF14 | Ring finger protein 14 | 0.999 |
| Testosterone | Testosterone | 0.999 |
| Bicalutamide | Bicalutamide | 0.999 |
| TMPRSS2 | transmembrane protease, serine 2 | 0.999 |

### 2.2 Molecular Docking

#### 2.2.1 Ligand preparation

The lowest energy 3D structures with corrected chiralities were produced by ligand optimization using the tool LigPrep tool. OPLS 2005 force field used and the process performed at neutral pH.

#### 2.2.2 Protein preparation and grid generation

Before the docking study, the biological unit of protein subjected to the protein preparation where it added with missing side chains and missing residues by the Prime tool. Further refining of the protein done by heavy atom and water molecule removal in restrained optimization. After the protein preparation, the receptor grid was generated using the OPLS 2005. The cubic box with specific dimensions generated around the centroid of the active site.

#### 2.2.3 Ligand Docking

The standard precision (SP) and extra precision (XP) flexible glide docking were used to screen the analogs, by using the Glide tool. For ligand atoms, Van der Waals factor and partial charge cut off were selected to be 0.80 and 0.15 respectively and ligands that are optimized used for this purpose.

#### 2.2.4 Compounds used for the study

1. Sesamol (SM)
2. 3′,4′-(Methylenedioxy) acetophenone (3’ MA)
3. Piperonly amine
4. 3,4-(Methylenedioxy) mandelic acid
5. Piperonylic acid
6. 3,4-(Methylenedioxy)aniline
7. Homopiperonylic acid
8. Piperonly alcohol
9. 3,4-(Methylenedioxy)-cinnamic acid

### 2.3 *In vitro* evaluation

#### 2.3.1 Evaluation of Cell Viability by MTT Assay

The cytotoxicity profile of SM and its derivatives on LNCaP, PC-3, and DU-145 cell lines was assessed by 3-(4,5-Dimethylthiazol-2-Yl)-2,5-Diphenyltetrazolium Bromide (MTT) assay. The 96-well plate was seeded with 4.0×10^3^ cells/well. SM and its derivatives at different concentrations from 156.25 μM −10 mM in 0.5% DMSO and vehicle control were incubated for 48 h. After incubation 20 μL MTT (2 mg/mL) is added and incubated at 37°C for four h, the precipitated formazan crystals were dissolved in 100 μL DMSO, and the optical density (OD) was measured at 540 nm by ELISA plate reader (BioTek, USA). The percentage of cell viability was determined by the equation: % of cell survival = {(OD of the sample)/ (OD of control)} ×100. Data were analyzed using GraphPad Prism 5.0. for half inhibitory concentration (IC_50_) determination.

#### 2.3.2 Quantitative Polymerase Chain Reaction (q-PCR) analysis

RNA expression studies of these lead compounds were further evaluated by q-PCR for the androgen signaling pathway. Androgen signaling genes like AR, FKBP5, PSA, and TMPRSS2 were evaluated. The LNCaP cells were treated with SM and 3’ MA at different concentrations for 48 hrs and compared with the standard nonsteroidal antiandrogen as enzalutamide (10 μM). Total RNA was extracted using TRIzol (Ambion, Waltham, MA) and reverse-transcribed using superscript II reverse transcriptase (Invitrogen). The qPCR study was carried out according to the established protocol using SYBR Green master mix (Applied Biosystems, Foster City, CA) on the Step One Plus (Applied Biosystems). The mRNA of interest showing expression was computed using the ΔΔCt technique and standardized to *GAPDH* (housekeeping gene) expression, and further relative expression was calculated and plotted.

Primers used in the study were as follows:

*AR* F – AATCCCACATCCTGCTCAAG R – GAGTCCAGGAGCTTGGTGAG
*FKBP5* F – GCAGGCGGTGATTCAGTAT
R – AGGTTCAGAAAGGCAGCAA
*PSA (KLK3)* F – GTCTGCGGCGGTGTTCTG
R – TGCCGACCCAGCAAGATC
*TMPRSS2* F – CAGGAGTGTACGGGAATGTGATGGT R – GATTAGCCGTCTGCCCTCATTTGT

#### 2.3.3 Western blot analysis

LNCaP cells were treated with different concentrations of sesamol and 3’MA (500, 1000 and 2000 μM) which was further compared with the standard AR antagonist enzalutamide (10 μM). The lysis buffer, RIPA (radioimmunoprecipitation assay) along with complete proteinase (Roche, Basel, Switzerland) and phosphatase inhibitors mixture (Calbiochem, Darmstadt, Germany) was used to prepare whole cell lysates. The experiment of western blot analysis involved the use of antibodies specific for androgen signaling and β-actin as housekeeping antibody (Cell signaling). The standard system of enhanced chemiluminescence was used to visualize the signals; this step was performed as per the manufacturer (GE Healthcare, Little Chalfont, UK) details. AR, PSA and Actin antibody from CST: 1:1000 diluted anti-AR (CST, 5153), 1:2000), 1:2000 diluted anti-PSA (CST, 5877), 1:5000 diluted anti-β-Actin (Abcam, ab6276) were used.

### 2.4 *In-vivo* Model

Testosterone undecanoate (TU) (Cernos Depot 1Gm Injection) was purchased from Sun Pharmaceutical. Sesamol and 3′,4′-(Methylenedioxy) acetophenone were purchased from Sigma-Aldrich. All the analytical grade chemicals were procured from Merck Limited, India and Hi-media. Plastic labware and syringes were procured from tarson and BD biosciences respectively. Rat PSA Prostate Specific Antigen (PSA) ELISA Kit (Catalog number: E-EL-R0796) is purchased from Elabscience, Houston, USA.

#### 2.4.1 Animals grouping and experimental design

The experimental design along with the grouping of animals are described in table 1.

#### 2.4.2 Induction of PCa in Rats by MNU and TU and the Effect of SM and 3’ MA on it

The study was conducted in compliance with Committee for the Purpose of Control and Supervision of Experiments on Animals (CPCSEA) guidelines with the prior approval of the Institutional Animal Ethical Committee at Central Animal Research Facility, Manipal) [Approved reference number - IAEC/KMC/04/2016]. Adult Male Sprague Dawley rats (180-200 g) were used for conducting the experiments. The experimental design is described in figure 1 was followed for the study.[18]

**Figure 1:**
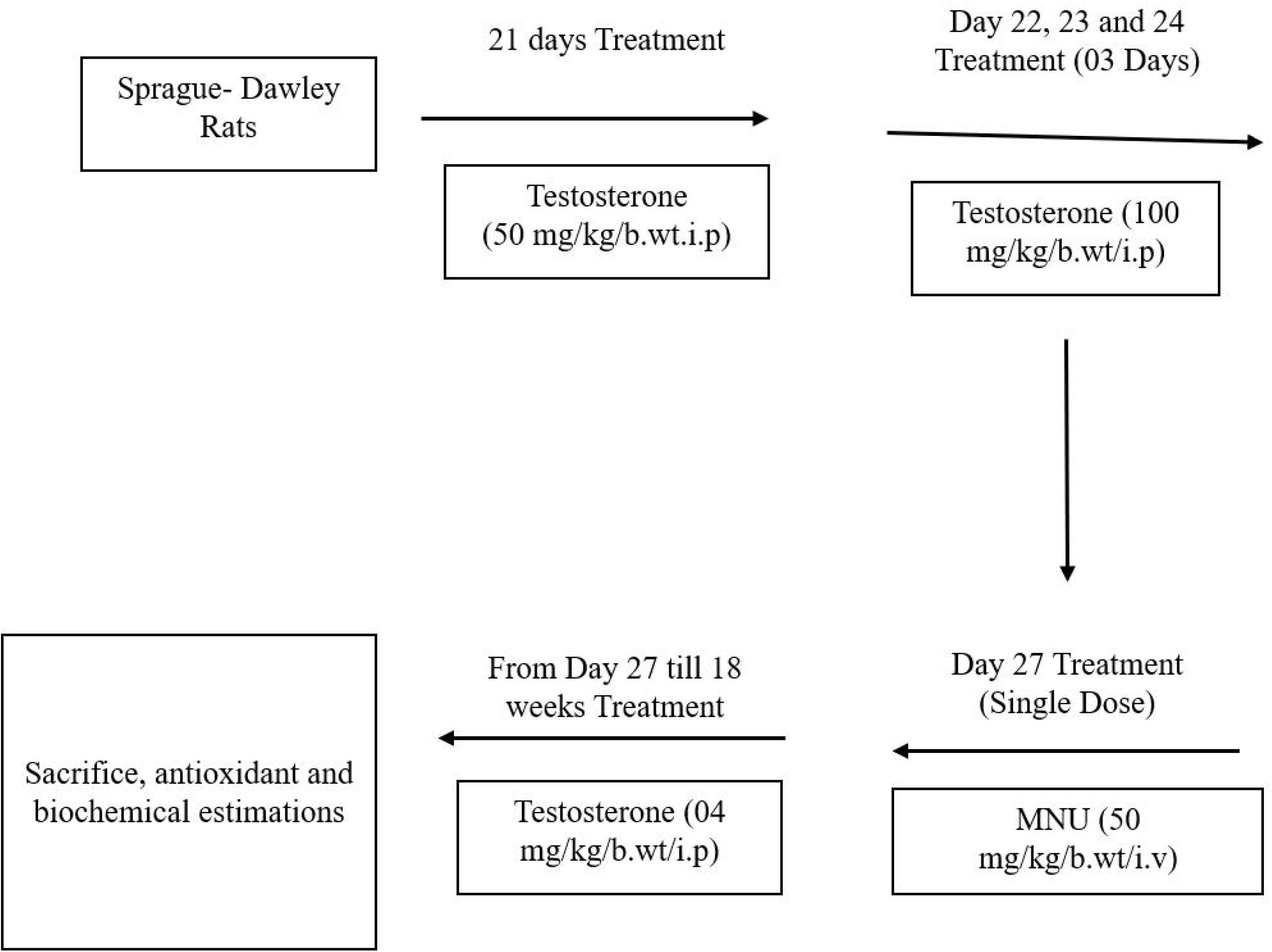
Experimental Design and dose regimen for TU and MNU to induce PCa

#### 2.4.3 Prostate weight

Prostate tissues were harvested, and weight was recorded.

#### 2.4.4 Histopathological examination

The prostate tissues were fixed and embedded in paraffin wax followed by sectioning. The thin sections were stained using hematoxylin and eosin. The stained sections were mounted on a glass slide and are observed under an inverted microscope and images were taken.

#### 2.4.5 Determination of antioxidant parameters

Antioxidant assays of ventral prostate homogenate {Catalase, Lipid Peroxidation, Nitrite, GSH} were screened.^[19]^

#### 2.4.6 Prostate homogenate preparation

Ventral prostate tissue was homogenized in ice-cold 0.1 mol/l of Tris- HCl buffer, pH 7.4, and centrifuged at 3000 rpm for 15 min. The supernatant was collected and used for antioxidant assay estimations. [18]

#### 2.4.7 Rat plasma prostate-specific antigen (PSA)estimation by ELISA method

The quantitative determination of rat PSA concentrations in rat plasma based on Sandwich-ELISA principle. Precoated plates specific to Rat PSA were used. Sample/standard added to the plate combined with the antibody. The plate is further treated with biotinylated detection antibody and Avidin-Horseradish Peroxidase (HRP) conjugate and incubated. The unbound components are washed away. The substrate solution is added to each well. Only in the presence of rat PSA, biotinylated detection antibody and Avidin-HRP conjugate will appear blue. The enzyme-substrate reaction is terminated by the addition of stop solution, and the color turns yellow and is measured at 450 nm ± 2 nm optical density (OD). The concentration of rat PSA in the samples was calculated from the standard calibration curve of PSA.

### 2.5 Statistical Analysis

The data were analyzed statistically and expressed as the mean ± SEM(n=6/3). Groups were compared using ANOVA followed by Tukey’s test for multiple comparisons. The level of significance set at p < 0.05.

## 3 Results

### 3.1 Stitch analysis

The network shows the correlation (Figure 2.I) of sesamol and its interactions with androgen receptor-regulated genes. Sesamol is lignan in nature; may shows phytoestrogen activity. Sesamol interacts with the Androgen receptor and regulates the androgen-regulated genes. The interaction of sesamol with AR genes, PSA (KLK3), TMPRSS2 indicates the possibility of the androgen-mediated pathway.

**Figure 2:**
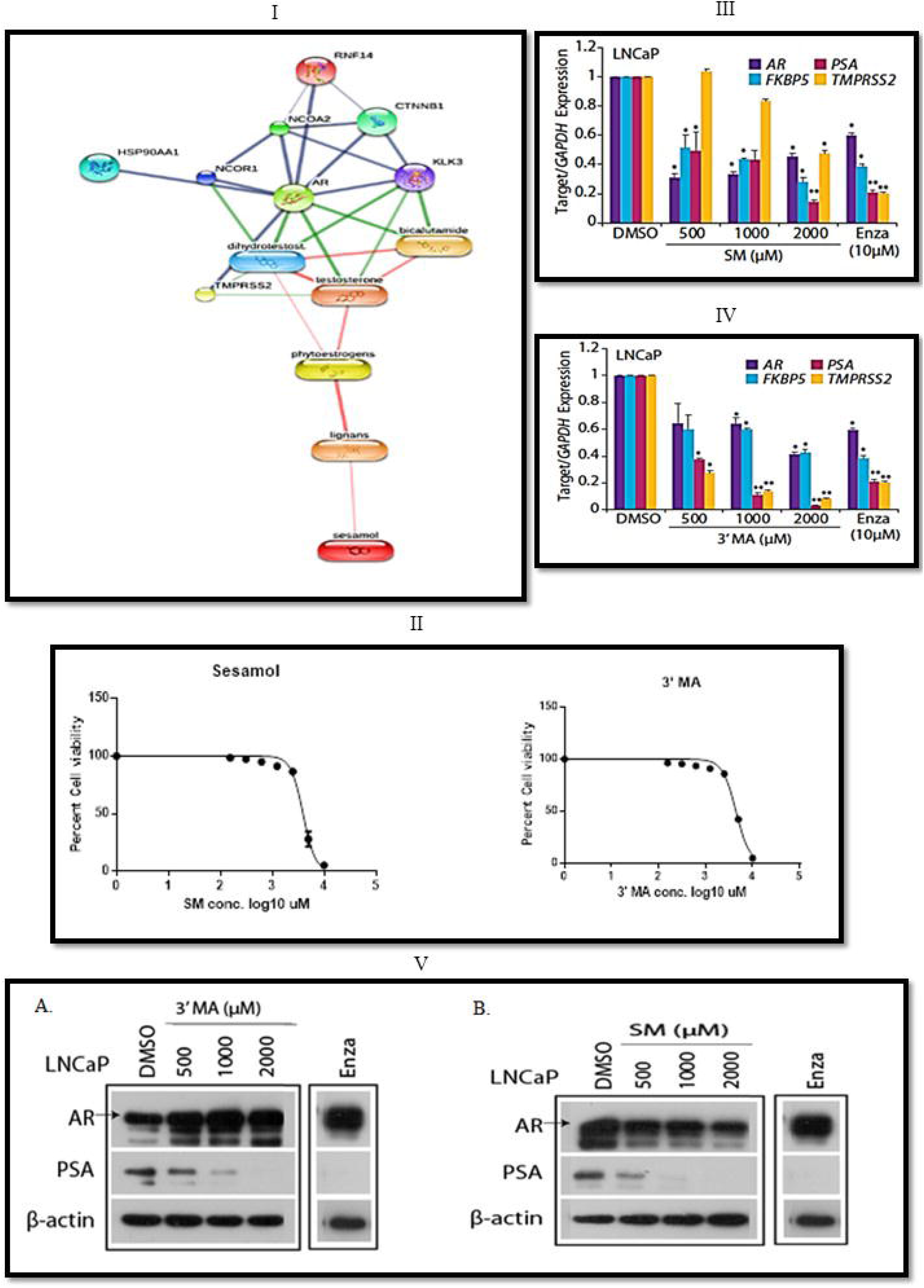
I. Network around Sesamol and Androgen-regulated genes; II. Dose-response curve of Sesamol and its derivatives on LNCaP cells; III. Effect of the Sesamol on AR-dependent gene transcription; IV. Effect of the 3’MA on AR-dependent gene transcription; V.A. Effect of the 3’MA on AR-dependent gene transcription; V.B. Effect of the SM on AR-dependent gene transcription

### 3.2 Schrodinger based Molecular docking for SM, and its derivatives

For the rational structure-based drug design, Sesamol and its eight different derivatives were investigated their binding affinity against AR. The human androgen receptor was downloaded from the Protein Data Bank using PDBID - 1E3G and 2AM9 respectively and docked using the Maestro Molecular Modelling platform by Schrodinger, LLC. The Dock score is represented in table 4 with a comparison of androgen receptors type.

**Table 4:** SP and XP docking score for SM and it’s derivatives

| Sr.<br>N. | Ligands | 1E3G |  | 2AM9 |  |
| --- | --- | --- | --- | --- | --- |
|  |  | SP<br>score | XP<br>Score | SP Score | XP<br>Score |
| **1** | Homopiperonylic acid | -7.527 | -9.823 | -6.986 | -6.544 |
| **2** | 3',4'-(Methylenedioxy) acetophenone | -7.7 | -7.188 | -6.965 | -6.49 |
| **3** | 3,4-(Methylenedioxy) mandelic acid | -7.65 | -8.129 | -6.443 | -7.188 |
| **4** | Piperonylic acid | -7.317 | -8.61 | -6.553 | -6.583 |
| **5** | Piperonly alcohol | -8.268 | -7.828 | -7.754 | -7.31 |
| **6** | Sesamol | -7.353 | -7.089 | -7.297 | -6.174 |
| **7** | 3,4-(Methylendioxy)-cinnamic | -7.822 | -8.809 | -5.981 | -6.857 |
| **8** | 3,4-(Methylenedioxy) aniline | -7.045 | -6.921 | -4.061 | -5.347 |
| **9** | Piperonly amine | -7.036 | -5.654 | -6.428 | -5.621 |

### 3.3 *In vitro* evaluation

#### 3.3.1 Cytotoxicity Assay

The compounds SM and 3’ MA showed the least IC_50_ (mM) as 3.94 and 4.43 (Figure 2.II) respectively on the other hand rest compounds were found to have very high IC_50_ values (Table 5).

**Table 5.** IC_50_ Concentrations of other Sesamol Derivatives on LNCaP cells

| Sr. No. | Name of Drugs | Mean IC<sub>50</sub> (mM) ±<br>Std. Error of Mean (SEM) |
| --- | --- | --- |
| 1 | Piperonyl amine | 12.214 ± 5.308 |
| 2 | 3,4-(Methylenedioxy) mandelic acid | 12.583 ± 4.367 |
| 3 | Piperonylic acid | 71.246 ± 0.638 |
| 4 | 3,4-(Methylenedioxy)aniline | 80.163 ± 1.797 |
| 5 | Homopiperonylic acid | 16.7 ± 2.980 |
| 6 | Piperonyl alcohol | 84.758 ± 8.991 |
| 7 | 3,4-(Methylenedioxy)-cinnamic acid | 78.632 ± 7.025 |

The IC_50_ (mM) in the androgen negative cell line was found to be more than 10 mM in PC-3 and DU-145 for the nine compounds. The values obtained indicated the drugs may be acting through the androgen signaling pathway so the androgen-independent cell lines were unaffected. The Vero cell lines, normal cells were also treated with these compounds and found to be safe in this assay; IC_50_ was more than 15 mM in each case.

#### 3.3.2 qPCR Studies

The experiment showed positive results as the expression was reduced by the SM and 3’ MA in all the genes at a higher dose. PSA notably was reduced by both in dose-dependent manners (Figure 2.III and Figure 2.IV). Thus, this confirmed the activity as anti-androgenic at the RNA level, and further protein expressions were confirmed by western blotting.

#### 3.3.3 Western blot analysis

The protein estimations of PSA in both cases were found to be dose-dependent inhibition in figure 2.V.A and 2.V.B.

### 3.4 *In-vivo* Model

#### 3.4.1 Prostate Weight

Animals treated with TU and MNU (disease control group) showed a significant increase in prostate weight of 78.68% compared to the normal control group.

In comparison with the disease control group, SM treatment (50 and 100 mg/kg/day, concomitantly administered with TU and MNU) significantly decreased the prostate weight by 25.14 % and 32.93%. Similarly, in the case of 3’ MA, at the dose of 50 and 100 mg/kg, there is a significant decrease in prostate weight as 31.43% and 57.44% which is comparable to the standard drug, finasteride (% reduction is 60.65) (Figure 3.I)

**Figure 3:**
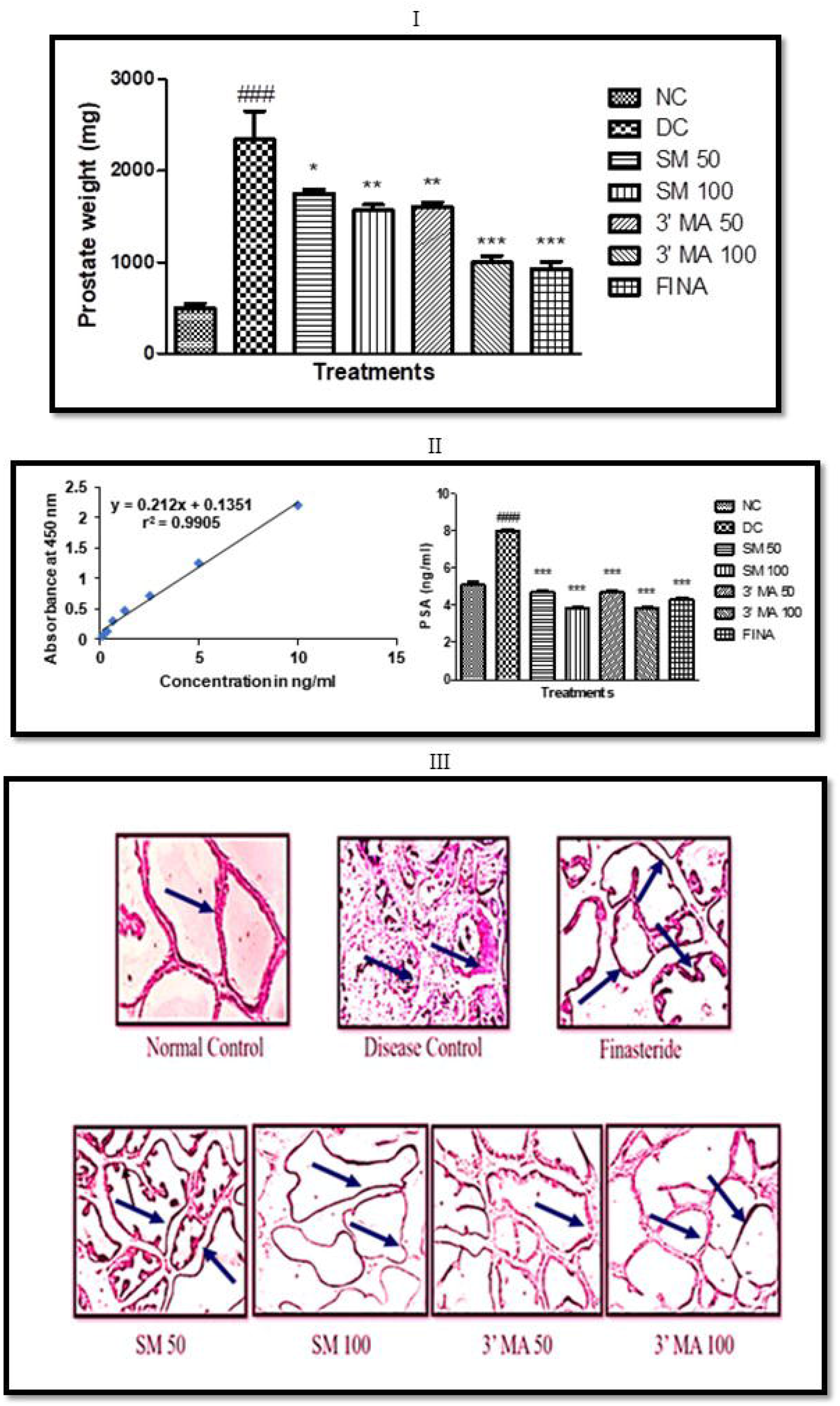
I. Effect of SM and 3’MA treatment on prostate weight; II. Effect of SM and 3’MA treatment on Prostate-Specific Antigen level in Rat Plasma; III-Histological examination of rat ventral prostates (100x images) – The arrow highlights the uniformity and distortion of the epithelial lining of cells

#### 3.4.2 Effect of SM and 3’MA treatment on antioxidant status of ventral prostate

Effect of SM and 3’MA treatment in two different doses (50 and 100 mg/kg, orally) on- antioxidant markers as nitrite, lipid peroxidation, catalase, and GSH level were evaluated and quantified as mentioned in Table 6.

**Table 6:** Quantitative estimation of Antioxidant parameters in prostate tissue homogenate

| **Animal Groups** | **Nitrite (nmol of CAT/mg protein)** | **CAT (nmol of CAT/mg protein)** | **Lipid peroxidation (nmol of MDA formed/mg protein)** | **GSH (nmol of GSH/mg protein)** |
| --- | --- | --- | --- | --- |
| NC | 2045±424.5 | 1.631±0.4895 | 102.7±20.88 | 1.77±0.3761 |
| DC | 4778±1575<sup>#</sup> | 0.4095±0.148<sup>ns</sup> | 232.2±48.21<sup>#</sup> | 0.8885±0.2928<sup>ns</sup> |
| SM 50 | 1805±459<sup>*</sup> | 2.975±0.4945<sup>ns</sup> | 47.52±3.956<sup>**</sup> | 2.816±0.6349<sup>**</sup> |
| SM 100 | 830±216.6<sup>**</sup> | 3.748±0.4511<sup>ns</sup> | 36.38±8.314<sup>***</sup> | 3.156±0.8108<sup>***</sup> |
| 3' MA 50 | 1081±777<sup>**</sup> | 3.153±0.6848<sup>ns</sup> | 71.81±16.11<sup>**</sup> | 1.912±0.5826<sup>**</sup> |
| 3' MA 100 | 385.6±248.5<sup>**</sup> | 3.123±0.1823<sup>ns</sup> | 76.54±46.88<sup>**</sup> | 2.569±1.194<sup>**</sup> |
| FINA | 1809±455.5<sup>*</sup> | 3.323±0.6438<sup>ns</sup> | 80.88±41<sup>**</sup> | 1.851±0.3344<sup>***</sup> |

Catalase and glutathione (GSH) level enhanced significantly in SM and 3’MA treatment group in a dose-dependent manner. While Nitrite and MDA level decreased in SM and 3’MA treated groups signifies the antioxidant nature. This antioxidant potential may help in reducing prostate cancer disease progression.

#### 3.4.3 Prostate Specific Antigen Estimation

The measurement of prostate-specific antigen in serum is an early detection marker in PCa. A significant decrease in PSA level (Figure 3.II) signifies, the influence of androgen signaling mediated pathway in SM and 3’MA treated groups.

#### 3.4.4 Histopathology – Prostate

The histopathological evaluation indicated marked changes in the prostatic histoarchitecture in SM and 3’MA treated groups and indicated protection in PCa (Figure 3.III).

## 4 Discussion

STITCH analysis indicated that SM may interact with androgen-regulated genes and can be used as an antiandrogen agent. The comparable docking score indicated the interaction of SM and its derivatives with AR receptors. MTT assay gives an insight on the SM and its one of the derivatives as 3′,4′-(Methylenedioxy) acetophenone (3’-MA) showing least IC_50_ as compared to the other derivatives. IC_50_ of SM and 3’ MA is 3940 and 4430 μM proposed to have the capacity to inhibits the growth of AR-positive human prostate cancer cell lines. To assess the antiandrogenic activity for SM and 3’MA, we evaluated their action on the fold expression change of the genes as androgen receptor gene (AR), FKBP5 (FKBP Prolyl Isomerase 5), Prostate-specific antigen (PSA/KLK3), Transmembrane Serine Protease 2 (TMPRSS2). in androgen-dependent LNCaP cancer cells (Figure 2.III and 2.IV). LNCaP cells have efficient receptors that are endogenous receptors of androgens. The mRNA expressions of AR, PSA, TMPRSS2, and FKBP5 which are tumor specific markers for PCa were significantly reduced by the treatments with SM and 3’MA.

The variation in PSA protein expression was also evaluated using Western blot analysis. It was observed that PSA expression levels were reduced by treatment with SM and 3’MA in a dose-dependent fashion in LNCaP cells. SM and 3’MA decreased the expression of PSA protein when compared with the standard AR antagonist enzalutamide (10 μM).SM is found to interact with AR and shown to cause AR ubiquitination with an alternate mechanism. 3’MA enhanced the protein level of the AR. Taken together, these results suggest that SM and 3’MA shown to interact with AR action in PCa cells and inhibition of AR regulated genes expression in the same manner as other natural AR antagonists reported in the literature.[20] Identification of AR-regulated genes and molecular pathway correlated with the PCa is of particular interest.

Relative prostate weight is used to evaluate the growth or development of PCa in rats. A rising in prostate gland weight (Disease Control) explained the effect of MNU induced a toxic effect on the prostate gland as compared to the normal control group. SM and 3’MA shown a dose-dependant decrease in prostate weight.

In antioxidant studies, SM and 3’MA treatment inhibited the level of malondialdehyde (MDA) and H_2_O_2_ in lipid peroxidation and catalase assay. The effect may be due to its scavenging properties of free radicals and thereby gave an insight into the protective mechanism. Glutathione (GSH) levels increased significantly (p<0.05) in the groups fed with SM and 3’MA while Nitrite content was decreased. The antioxidant potential of SM and 3’MA might help in the prevention of prostate cancer progression. To confirm these results, we have examined the rat plasma PSA level and histopathological alterations in the ventral prostate section.

PSA is synthesized in the ductal and acinar epithelium of the prostate gland and secreted into the seminal plasma. The function of PSA is to liquefy the seminal coagulum by proteolysis, with the release of the entrapped spermatozoa and may support fertilization. [20] PSA’s rapid rise is closely associated with prostate cancer. PSA is typically secreted as pro-PSA and activated outside the normal or malignant epithelial cell. In normal cells, the pro-PSA peptide is correctly removed to generate enzymatically active PSA. In cancers, the pro-PSA peptide is aberrantly processed to produce a series of isoforms that can be measured in the blood.

Therefore, plasma PSA concentrations in rats were determined to evaluate the effect of SM and 3’MA in prostate cancer. PSA showed a significant increase in the disease control group regarding normal control. This may be due to disruption of the epithelium leads to diffusion of PSA into the blood. Histological study in the rats, SM and 3’MA orally administered showed reduced proliferative lesions and cell injuries and considerable improvement in the prostatic histoarchitecture in SM and 3’MA treated groups. The data obtained from the study correlate the protective mechanism of SM and 3’MA in the treatment of PCa.

## 5 Conclusion

Results from the studies reveal that SM and 3’MA could be hopeful molecules for further assessment for PCa treatment option for preventive therapy. In the present investigation, sesamol and its derivative (3’-MA) prevented the development of PCa by interfering androgen-regulated pathway.

## Acknowledgements

The authors thank the Department of Pharmacognosy, Department of Pharmacology and Manipal - Schrodinger Centre for Molecular Simulations, Manipal College of Pharmaceutical Sciences and Manipal Academy of Higher Education, Manipal, for providing all the infrastructural facilities to carry out the research work. The support provided by Dr. Bushra Ateeq, Molecular Oncology Laboratory, Department of BSBE, IIT-Kanpur for this work is grateful. Her student Mr. Vipul Bhatia helped us to perform some experiments, a great thanks to both.

